# Targeted cancer cell killing by highly selective miRNA-triggered activation of a prokaryotic toxin-antitoxin system

**DOI:** 10.1101/695114

**Authors:** Alice Turnbull, Camino Bermejo-Rodríguez, Mark A. Preston, María Garrido-Barros, Belén Pimentel, Guillermo de la Cueva-Méndez

## Abstract

Although not evolved to function in eukaryotes, prokaryotic toxin Kid induces apoptosis in human cells, and this is avoided by co-expression of its neutralizing antitoxin, Kis. Inspired by the way Kid becomes active in bacterial cells we had previously engineered a synthetic toxin-antitoxin system bearing a Kis protein variant that is selectively degraded in cells expressing viral oncoprotein E6, thus achieving highly selective killing of cancer cells transformed by human papillomavirus. Here we aimed to broaden the type of oncogenic insults, and therefore of cancer cells, that can be targeted using this approach. We show that appropriate linkage of the *kis* gene to a single, fully complementary, target site for an oncogenic human microRNA enables the construction of a synthetic toxin-antitoxin pair that selectively kills cancer cells overexpressing that particular microRNA. Importantly, the resulting system spares non-targeted cells from collateral damage, even when they overexpress highly homologous, though non-targeted, microRNAs.

## Introduction

Prokaryotic organisms and many of their resident plasmids have evolved toxin-antitoxin (TA) systems that function as stress response elements. These encode intracellular toxins that pass unnoticed to host cells due to the neutralizing action of co-expressed antitoxins. However, the latter are rapidly degraded following cellular exposure to certain types of stress, liberating free toxin. Some of these toxins trigger selective degradation of mRNAs, which reversibly inhibits cell growth and alters gene expression in ways that help arrested cells to effectively respond to, and survive, the inducing stress.^1–3^

For instance, the Kid-Kis TA pair functions as a prokaryotic plasmid rescue system. Bacterial cells carrying plasmid R1 produce toxin Kid and neutralizing amounts of antitoxin Kis, and thus proliferate unhindered. However, this only happens when host cells contain enough R1 copies to ensure that the plasmid will be transmitted to both daughter cells on cell division. When R1 copies are insufficient to guarantee this, proteolytic degradation of Kis frees Kid, and the toxin targets mRNAs at 5’-UUACU-3’ sites, destroying them. This not only inhibits cell division, which avoids production of plasmid-free cells, but also increases R1 replication rates. As a consequence, plasmid copy numbers raise rapidly in arrested cells, eliminating the risk that they produce R1-free descendants once cell growth resumes. When copy number has recovered, Kid is re-neutralized by newly produced Kis, and cells are allowed to proliferate again.^4–7^

The Kid-Kis system has evolved to exquisitely support the maintenance of R1 in its natural prokaryotic environments, with toxin activation resulting in a reversible cytostasis, which is perfectly suited for R1 replication, copy number rescue and plasmid survival. In contrast, the expression of Kid triggers apoptosis in evolutionarily distant human cells, something that is prevented by co-expression of Kis.^8^ We proposed to exploit this effect, and use Kid and Kis to engineer synthetic TA systems able to achieve selective killing of cancer cells, taking advantage of the fact that many oncogenic insults reduce the intracellular concentration of specific proteins.^9–12^ Specifically, we produced Kis antitoxin variants that were degraded by the proteasome in cancer cells expressing human papillomavirus oncoprotein E6, thus leaving Kid free to trigger their death by apoptosis. Off-target delivery of this synthetic TA system to other cells had not effect on cell viability, as these cells could maintain neutralizing amounts of the antitoxin.^13^ These results suggested that our approach has potential to offer smart anticancer systems that avoid causing collateral damage to non-targeted cell, something that has so far proved very challenging to achieve in the clinic.^14–16^

In this work we wished to broaden the type of oncogenic insults, and therefore of cancer cells, that could be targeted using our strategy. Specifically, we aimed at engineering a new class of synthetic TA-pair that becomes active in human cells suffering microRNA-driven oncogenic stress. These small non-coding RNAs (miRNAs) inhibit the translation, or induce the degradation, of target mRNAs through complementary base pairing with target sites (miRts), most frequently located in their untranslated regions.^17,18^ The degree of base pairing seems to determine whether a given miRNA-miRts interaction causes translational inhibition (reduced pairing) or degradation (extended pairing) of targeted transcripts^18–19^, although more recent results show that some miRNAs can induce translational inhibition of target transcripts, instead of their degradation, in spite of the fact that they anneal the latter with full complementarity.^20^ Many studies have demonstrated that miRNAs are deregulated in cancer cells and can function as oncogenes. For instance, miR373 stimulates proliferation, migration and metastasis of cancer cells,^21–22^ and is overexpressed in many human cancers.^23–29^

## Results and Discussion

We hypothesized that a vector bearing independent transcriptional units for the expression of Kid and Kis in human cells, as well as a miRts for miR373 (miR373ts) downstream of the *kis* gene, would result in selective killing of cancer cells overexpressing this miRNA (Figure 1A). We noted that human cells express more than 2,500 different miRNAs, and that many of them share considerable sequence homology^30^ (Figure 1B), which could lead to inappropriate activation of Kid in non-targeted cells expressing miR373 homologues. To avoid this risk our design included a single miR373ts placed 6 nucleotides downstream of the *kis* stop codon, a location that is considered inhospitable for miRNAs that do not bind with full complementarity to their target sites, due to ribosomal interference^31,32^ (Figure 1C).

**Figure 1.**
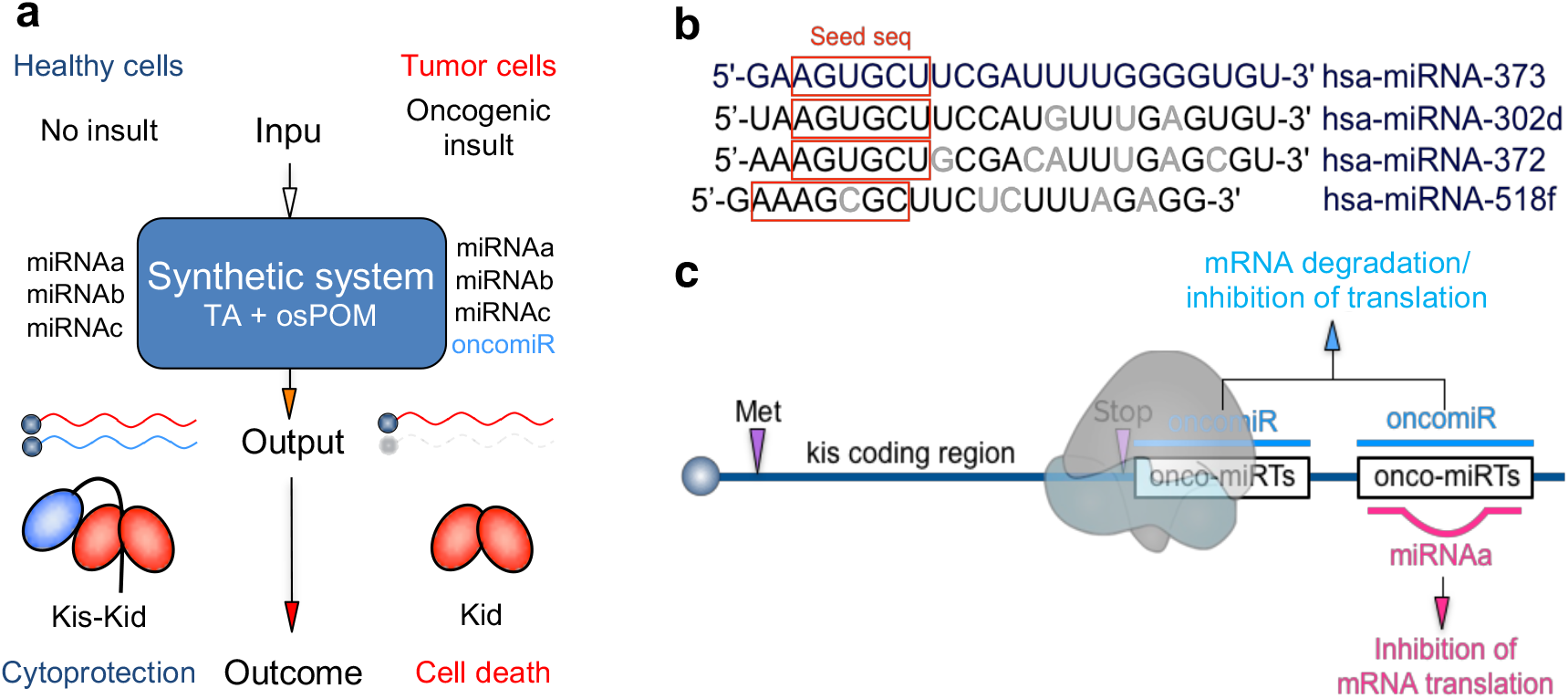
Rationale for the construction of a synthetic TA-based system that selectively kills human cells oncogenically stressed by miRNAs. (a) Diagram depicting the intended behavior of the synthetic system. Genes *kid* and *kis* are transcribed from separate eukaryotic transcriptional units in an appropriate expression vector. A DNA sequence 100% complementary to the targeted onco-miRNA is physically linked to the antitoxin gene and functions as an osPOM (i.e. onco-specific Protein Output Modifier) element. This enables the system to distinguish cells suffering the targeted miRNA-mediated oncogenic insult (input) from those that do not and, in response to this, induce the degradation of antitoxin transcripts in the former cells, increasing their intracellular Kid/Kis ratios (output), and inducing their death by apoptosis (outcome). (b) Sequence alignment of hsa-miR373 and its homologues hsa-miR302d, hsa-miR372 and hsa-miR518f. Seed sequences are boxed in red, and sequence differences between the latter miRNAs and reference hsa-miR373 are highlighted in grey. (c) Scheme of the rationale for the precise location of miRNA-responsive osPOMs in our system. Not fully complementary recognition of target sites by miRNAs can result in the inhibition of translation from affected transcripts. miRNA displacement by ribosomal interference imposes restrictions to miRNA target site recognition in the proximity of stop codons, limiting the effect to miRNAs that are fully complementary to its target sequence (blue miRNAs). Accordingly, placing a single miRNA target site immediately downstream of the *kis* stop codon in our vector should limit toxin activation to cells containing its fully complementary miRNA, thus avoiding serendipitous, off target, induction of apoptosis in cells containing any other miRNA (pink miRNA).

To test our hypothesis, we produced a cell line in which we could regulate the expression levels of miR373. We cloned the stem-loop pri-miR373 precursor downstream of a minimal CMV promoter regulated by doxycycline. The resulting plasmid (pTet-miR373; Figure 2A) was integrated in the genome of 293TRSID, a human cell line that allows both positive and negative regulation of the promoter above.

**Figure 2.**
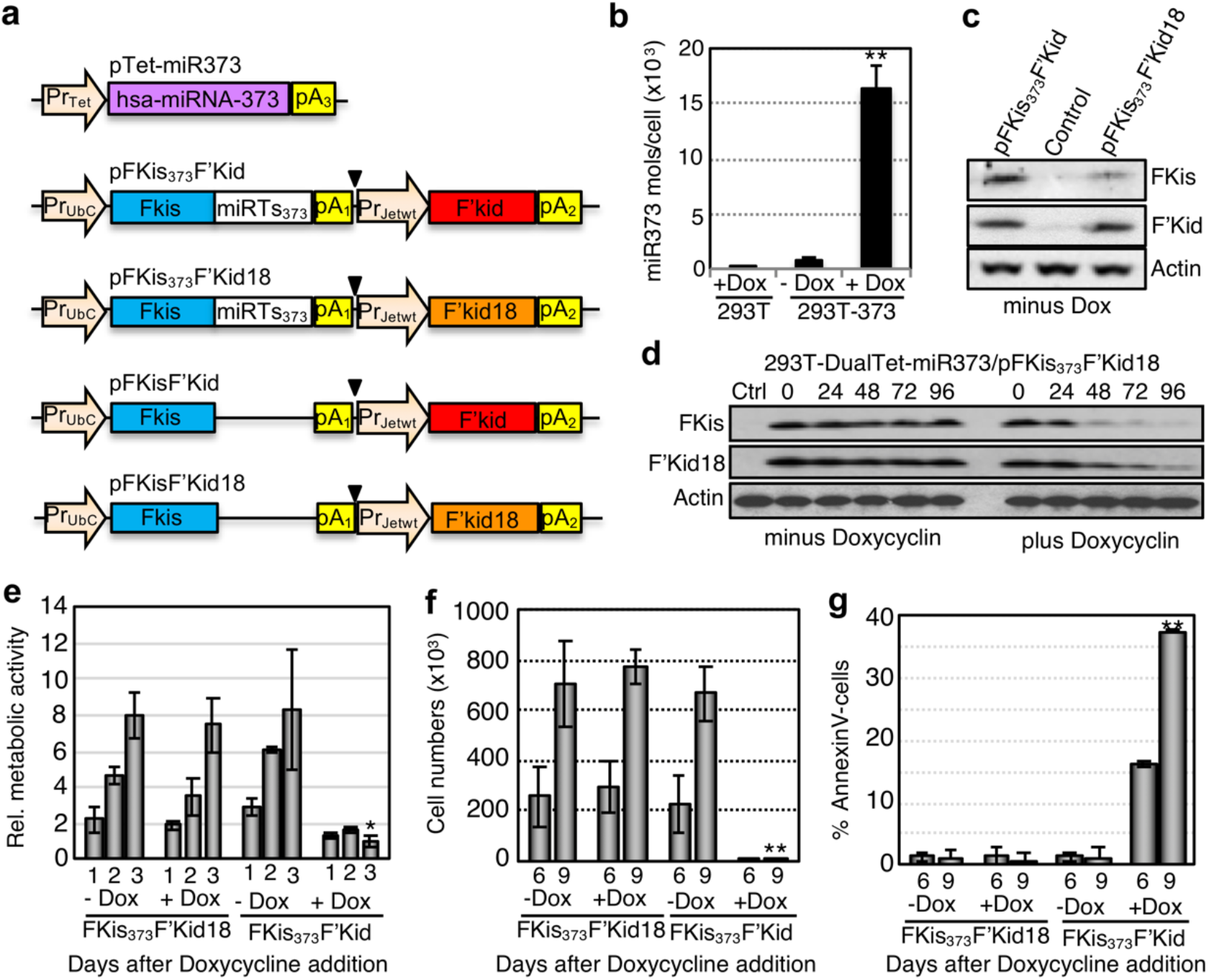
Functional characterization of the system using inducible miR373-expressing cell lines. (a) Scheme of the miR373-, Fkis- and F’kid-(or F’kid18-) bearing transcriptional units in all vectors used to produce our cell lines. (b) Average number of miR373 molecules per cell in 293TRSID-Dual-Tet-miR373 cell cultures grown in the presence (1μg/ml) or the absence of doxycycline, and in parental 293TRSID cell cultures (Ctrl) also grown in the presence of doxycycline. (c) Western blot using antibodies against the FLAG epitope and human actin, performed on protein extracts produced from 293TRSID-Dual-Tet-miR373-pFKis373F’Kid, 293TRSID-Dual-Tet-miR373-pFKis373F’Kid18, or their parental cells (control), cultured in the absence of doxycycline. (d) Western blot using an antibody against the FLAG epitope, and performed on protein extracts produced from 293TRSID-Dual-Tet-miR373-pFKis373F’Kid18 cells cultured for the indicated hours in the absence or the presence of doxycycline (1μg/ml). Molecular weights for FKis, F’Kid and actin are 10.6, 13.15 and 42 kDa, respectively (e) Fold change of the average metabolic activity observed in 293TRSID-Dual-Tet-miR373-pFKis373F’Kid and 293TRSID-Dual-Tet-miR373-pFKis373F’Kid18 cells cultured for the indicated days in the absence or the presence of doxycycline (1μg/ml). (f) Average number of cells in 293TRSID-Dual-Tet-miR373-pFKis373F’Kid and 293TRSID-Dual-Tet-miR373-pFKis373F’Kid18 cells cultured for the indicated days in the absence or the presence of doxycycline (1μg/ml). (g) Average percentage of annexin-positive cells (a surrogate for apoptosis) in the samples analysed in (f). * and ** denote an statistically significant difference at 90% and 95% confidence interval, respectively, between +Dox and -Dox samples. Experiments were repeated three times, and all measurements were carried out in triplicates.

The resulting cell line (293TRSID-Dual-Tet-miR373) expressed low levels of miR373 in the absence of doxycycline, but very high it its presence (Figure 2B). We also made plasmids allowing the independent co-expression of Flag tagged-Kis and -Kid (or its inactive mutant, Kid18) proteins in human cells, both bearing or lacking the perfect miR373ts downstream of the *kis* gene (pFKisF’Kid, pFKisF’Kid18, pFKis_373_F’Kid and pFKis_373_F’Kid18; Figure 2A and Figure 1 in supporting information). We integrated these plasmids in the genome of 293TRSID-Dual-Tet-miR373 cells, and selected clones that expressed similar levels of F’Kid and F’Kid18 (Figure 2C). Next, we grew the stable pFKis_373_F’Kid18 clone in the presence and the absence of doxycycline and compared the amount of FKis and F’Kid18 produced by these samples over time. This showed that in the absence of doxycycline the amounts of FKis and F’Kid18 remained unchanged over time. However, addition of doxycycline and induction of miR373 transcription led to a rapid reduction of FKis expression (Figures 2D and supplementary Fig. 2). The latter was followed by a progressive reduction in the amounts of F’Kid18, indicating that, in human cells, the toxin is less stable in its free form than when bound to the antitoxin.

To analyse whether these changes were sufficient to enable the activation of Kid toxicity, we followed net metabolic activities (a surrogate for cell proliferation/viability) in FKis_373_F’Kid18- and FKis_373_F’Kid-cultures grown in the absence and the presence of doxycycline. Activities increased at similar rates in the case of FKis_373_F’Kid18-cultured in both conditions, and of FKis_373_F’Kid-cells grown in the absence of doxycycline. However, they did not change, and always remained very low, in the case FKis_373_F’Kid-cells grown in the presence of doxycycline (Figure 2E). We also measured the total number of cells and the percentage of apoptotic cells in the samples above, 6 and 9 days after the addition of doxycycline. These experiments showed a consistent increase in cell numbers in FKis_373_F’Kid18-cultures grown in the presence and the absence of doxycycline, and in FKis_373_F’Kid-cultures grown in the absence of doxycycline (pictures of these cells are shown in supplementary Fig. 3). However, when doxycycline was added to FKis_373_F’Kid-cultures, cell proliferation was severely inhibited (Figure 2F) and the percentage of apoptotic cells greatly increased (Figure 2G). Altogether, these results demonstrated that our vector ensures sufficient expression of antitoxin Kis to neutralize co-expressed Kid in cells containing low amounts of miR373. They also showed that overproduction of miR373 (i.e. which also takes place in some cancer cells), largely reduces the amount of antitoxin protein that these cells can produce from the vector, thus increasing intracellular Kid/Kis ratios and triggering apoptotic cell death.

We next tested whether our design would avoid unintended activation of the toxin by miRNAs sharing sequence homology with miR373 (Figure 1B). We transfected HEK293T cells with pFKisF’Kid18 or pFKis_373_F’Kid18, alone or mixed with either miR373-, or its homologues miR372-(69.56% identity; same seed match), miR302d-(78.26% identity; same seed match), miR518f-(71.43% identity; imperfect seed match), or miR-artif-(artificial miRNA mimic with imperfect seed match, but otherwise perfect sequence identity; 86%) mimics. We analyzed protein extracts from all these samples 24 hours after transfection by western blot, using an anti-FLAG antibody (Figure 3A). Expression of FKis was not reduced by any of the miR373 homologues compared to the control (no miRNA) sample. Reassuringly, all these miRNA-mimics were able to repress expression of FKis in cells co-transfected with pFKis_miRts_F’Kid18variants carrying their corresponding, fully complementary, miRts (Figure 3B). These results confirm the high selectivity of our system, which is able to differentially modulate Kid-to-Kis ratios in response to a predetermined miRNA, but not to other, highly homologous, ones.

**Figure 3.**
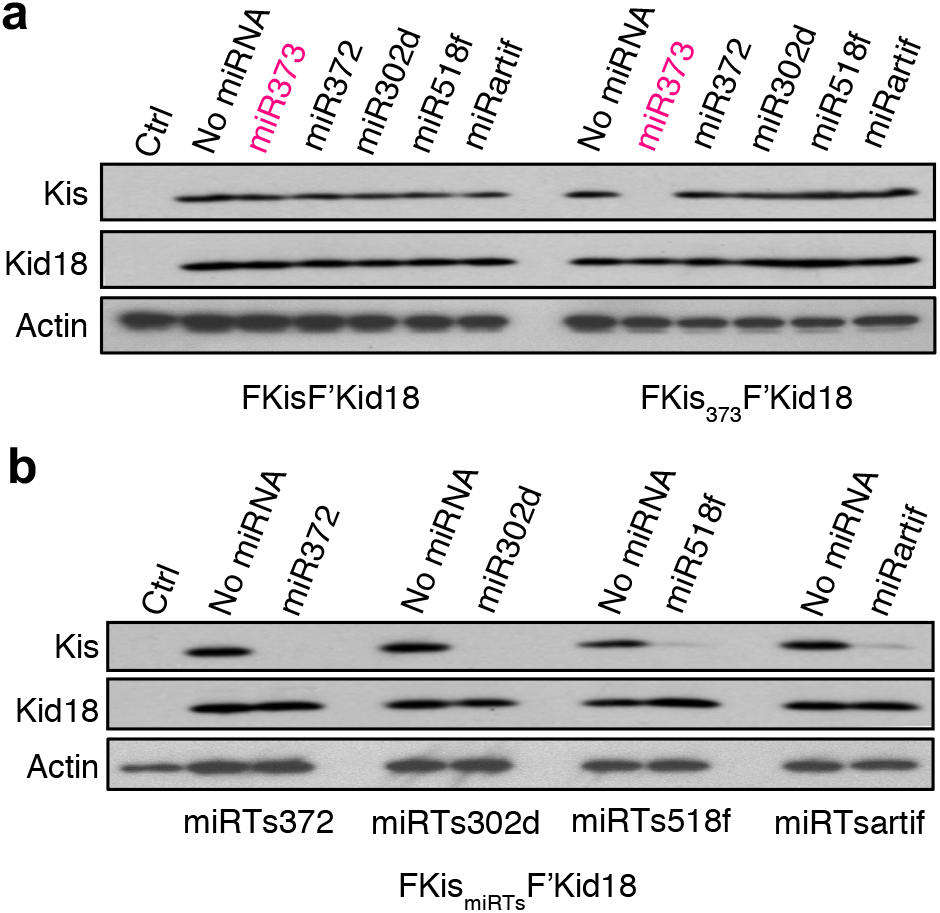
miRNA discriminatory selectivity of the synthetic TA-based cell killing system. (a) Western blot using an antibody against the FLAG epitope, and performed on protein extracts produced from HEK293T cells 24 h after being transfected with plasmids pFKisF’Kid18 or pFKis_373_F’Kid18, alone or mixed with 10 pmol of the indicated miRNA mimic. (b) Western blot using an antibody against the FLAG epitope, and performed on protein extracts produced from HEK293T cells 24 h after being transfected with plasmids pFKis_372_F’Kid18, pFKis_302_dF’Kid18, pFKis_518f_F’Kid18 or pFKis_artif_2F’Kid18 alone or mixed with 10 pmol of the indicated miRNA mimic. Ctrl denotes the use of extracts produced from untransfected HEK293T cells.

We grew the 293T-DualTet-miR373-FKis373F’Kid cells (used in Fig2) in the absence of doxycycline to avoid overexpression of the inducible miR373 integrated copy, and transfected them with 10 pmols of miR373, miR372, miR518f or miR302d. This experiment was designed to examine whether our system would achieve selective killing of cells transfected with miR373 only, and not with any of its other highly homologous miRNAs. For this, we analysed the net metabolic activity and the numbers of total and dead cells in these transfected cultures over time (supplementary Fig. 4). Our results confirmed a severe inhibition of the net metabolic activity and cell proliferation in cells transfected with miR373, which was accompanied by an increase of apoptotic cells in these cultures. None of these effects were observed with any of the other miRNAs, although we found that miR302d caused a slight inhibition of cell proliferation. However, this was neither paralleled by a reduction of net metabolic activity nor by an increase of apoptotic cell numbers (as expected in the presence of Kid activation) in this sample. The latter, and the observation that overexpression of miR302d in cancer cells induces their eventual arrest in the G1/S transition of the cell cycle,^33–35^ strongly suggested that this result was due to an intrinsic effect of miR302d function, and not to inhibition of Kis expression (as also confirmed by our results in Fig. 3A)

Finally, we tested the ability of our system to induce selective killing of tumor-derived cells that overexpress miR373. Extensive evidence describes the oncogenic character of this miRNA, and its upregulation has been confirmed in many human cancers, including a large number of testicular germ cell tumors (TGCTs).^22, 24–29^ Two TGCT cell lines expressing high (2102Ep) or low (PA-1) levels of miR373 (Figure 4A) were transfected with pFKis_373_F’Kid18 or pFKisF’Kid18, and we analyzed the expression of F’Kis and F’Kid18 in these samples 24 hours after transfection.

**Figure 4.**
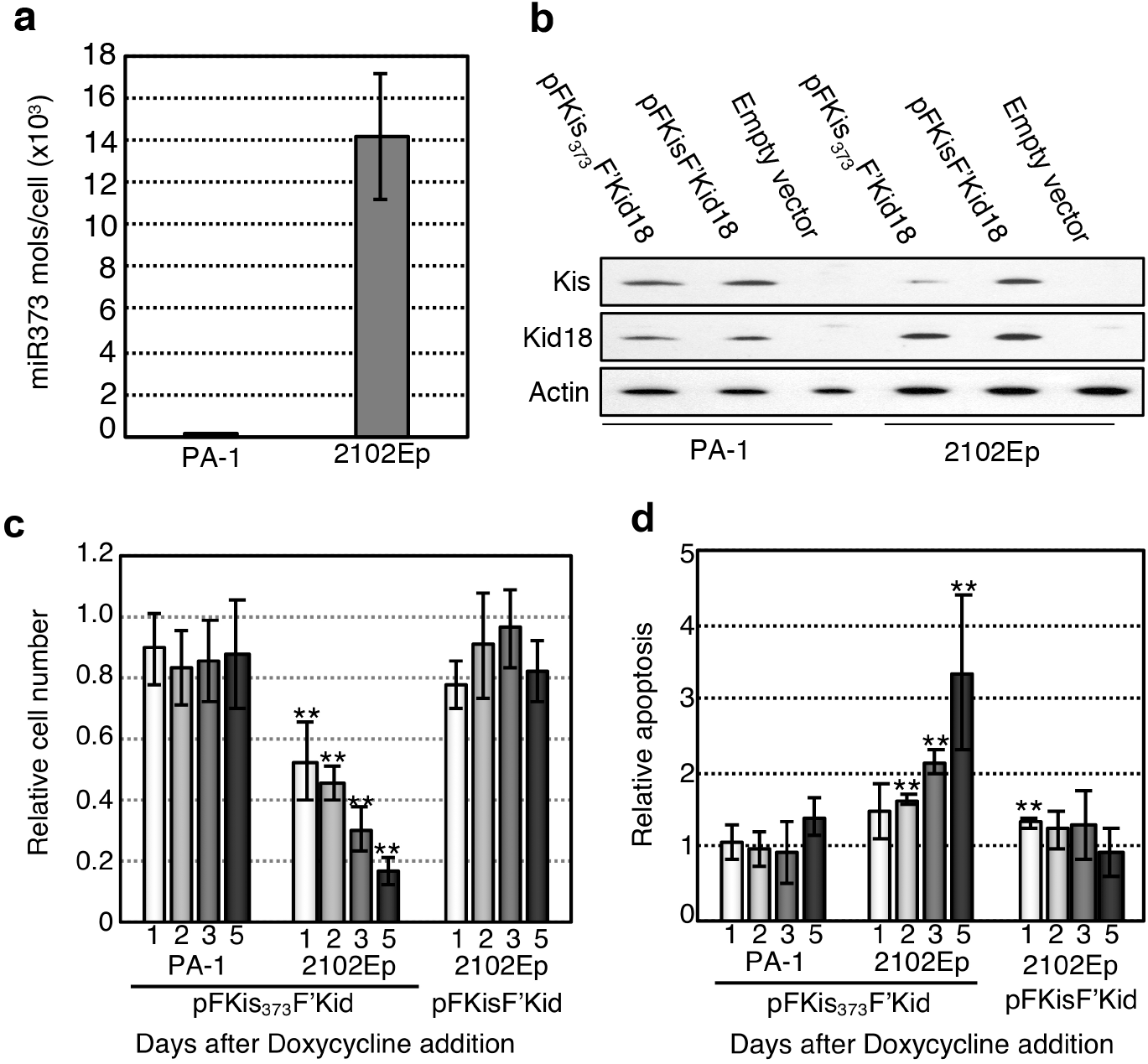
Functional characterization of the system using cancer cell lines. (a) Average number of hsa-miR373 molecules per cell in cultured Testicular Germ Cell Tumor (GCT) cell lines PA-1 and 2102Ep. (b) Western blot using antibodies against the FLAG epitope, performed on protein extracts produced from PA-1 and 2102Ep cells transfected with pFKisF’Kid18, pFKis_373_F’Kid18 or their empty parental vector. (c) Relative number of cells in cultures of PA-1 and 2102Ep cells transfected with pFKis_373_F’Kid of pFKisF’Kid for the indicated days. Values reflect ratios between the samples above and reference samples transfected with their control plasmids, pFKis_373_F’Kid18 of pFKisF’Kid18 (d) Average fold change in apoptotic cell numbers in the samples analysed in (c). ** denotes an statistically significant difference at 95% confidence interval, between Kid- and control Kid18-transfected samples. Experiments were repeated three times, and all measurements were carried out in triplicates.

Toxin-to-antitoxin ratios were very similar in 2102Ep and PA-1 cells transfected with pFKisF’Kid18, as well as in AP-1 cells transfected with pFKis_373_F’Kid18. However, in agreement with our earlier observations, the presence of miR373ts in this plasmid decreased FKis production (and therefore increased F’Kid18 to FKis ratios) considerably in 2102Ep cells (Figure 4B). Next, we measured cell proliferation and cell death rates in 2102Ep and AP-1 cells transfected with pFKis_373_F’Kid or control pFKisF’Kid, comparing them to those observed in the same cells transfected with control plasmids pFKis_373_F’Kid18 or pFKisF’Kid18. Whilst cell proliferation rates were similar for AP-1 cells transfected with pFKis_373_F’Kid and with control pFKis_373_F’Kid18, they decreased progressively in the case of 2102Ep cells transfected with pFKis_373_F’Kid, compared to their pFKis_373_F’Kid18 controls. This effect was entirely dependent on the presence of the miR373ts downstream of the *kis* gene, as it was not observed when the 2102Ep cells were transfected with control plasmid pFKisF’Kid (Figure 4C). In agreement with our earlier findings, inhibition of cell growth correlated with an increase in apoptotic cell numbers, which was not observed in control experiments (Figure 4D). These results further confirmed that our approach enabled highly selective killing of cancer cells exposed to predetermined miRNA-mediated oncogenic insults, sparing other cells from collateral, off-target, toxicity.

## Conclusions

Repurposing of functional elements enabling post-transcriptional regulation of protein outputs in human cells constitutes a useful strategy for the construction of smart therapeutic agents.^30–32.36^ Here, we have used the prokaryotic Kid-Kis TA pair to engineer a system capable of responding to abnormally high levels of a particular onco-miRNA in the complex intracellular environment of human cells and, in response to this stimulus, achieve selective activation of the system and cellular toxicity. In cancer cells overexpressing a specific onco-miRNA, our system interfaces with endogenous miRNA-dependent signaling pathways to inhibit expression of antitoxin *kis*-miRts. As a consequence, the default, cytoprotective ratio of Kid to Kis is disrupted, and apoptotic cell death is induced. Importantly, our system is extremely selective, avoiding accidental activation of the toxin in non-targeted cells, even if these overexpress miRNAs that are highly homologous to the one selected to trigger TA toxicity. This approach could help to develop personalized anticancer agents that are both cytotoxic and selective, and thereby avoid off-target effects in patients. In this context, we advocate the value of genetic circuits of the type presented here for addressing such a complex challenge, which has been hardly tractable by other means thus far.

## Material and Methods

### Oligonucleotides, plasmids, cell lines and transfection conditions

Oligonucleotides used in this work and a detailed description of the steps followed for the construction of plasmids and cell lines used in this work are shown in Supporting Information.

### Quantification of hsa-miR373 expression

Total miRNA was extracted from triplicate cultures (T75 flasks; 80% confluency), using Ambion’s miRVana kit, counting first the number of cell used in each preparation. Each qRT-PCR reaction was also carried out in triplicates, using the appropriate TaqMan miRNA assay kit and following manufacturer’s instructions. Relative quantification of miR373 was performed following a standard curve method using the RNU24 housekeeping gene for normalization. Absolute copy numbers of miR373 in each sample was calculated by interpolation with data obtained from parallel TaqMan reactions performed on samples containing increasing copy numbers of a miR373 mimic (ranging from 10^4^ to 10^10^) (Ambion; Table 1 in supporting information). The absolute number of miR373 copies per cell was calculated by dividing the miRNA copy number by the total number of cells used to prepare the miRNA sample.

### Analysis of FKis and F’Kid18 expression

For the experiment in Fig. 2C, 293TRSID-Dual-Tet-miR373 cells or their stable FKis_373_F’Kid- and FKis_373_F’Kid18-variants were cultured in the absence of doxycycline and harvested. Cell pellets were resuspended in 2 volumes of complete TNESV Protein extraction buffer (50 mM Tris-HCl (pH 7.4), 1% Nonidet P-40, 2 mM EDTA, 100 mM NaCl, 10 mM NaF, 1 mM Na3VO4, 1mM Pefabloc), previously supplemented with one tablet (per 10 ml of buffer volume) of complete protease inhibitor cocktail (Roche). Cell suspensions were incubated on ice for 15 min, vortexed every 5 min, and then centrifuged at 13000 rpm for 30 min at 4°C, to separate soluble proteins extracts from cellular debris. 20 μg of soluble protein per sample were run in a 4-20% Tris-Glycine gradient polyacrylamide gel (Invitrogen) and transferred to a ProTran NitroCellulose transfer membrane (Whatman) for detection by western blot, using a mouse monoclonal anti-FLAG antibody (Sigma), following manufacturer’s recommendations. For the experiment in Fig. 2D, we proceeded as above, but only using stable FKis_373_F’Kid18-cells grown with (1 μg/ml) or without doxycycline for the indicated timepoints. For the experiments in Fig. 3, we proceeded as above, but using HEK293T cells harvested 24h after being transiently transfected with the indicated pDNA:miRNA mimic mixtures. Finally, for the experiment in Fig. 4B, we followed the same protocol, but using 2102Ep and PA-1 cell cultures, 48 h after being transiently nucleofected with the indicated plasmids.

### Analysis of cell proliferation and cell death

For the experiment in Fig. 2E, 2×10^4^ 293TRSID-Dual-Tet-miR373 cells stably transfected with pFKis373F’Kid or with pFKis373F’Kid18 were seeded in triplicates and allowed to adhere for 24 h. We then determined the metabolic activity in our samples using Cell Counting Kit 8(CCK-8; Dojinjo), a colorimetric assay that quantifies the amount of NADH and NADPH (a surrogate for metabolic activity, and therefore for cell viability and cell numbers) produced by cultured cells, following manufacturer’s instructions. Measurements were carried out in cultures grown in the absence or the presence (1 μg/ml) of doxycycline, and immediately before (day 0), or 1, 2 and 3 days after addition of doxycycline to the corresponding samples. Experiments were carried out in triplicates, and data plotted as fold change in metabolic activity with respect to that observed on day 0 for each sample. For experiments in Figures 2F and 2G, 10^4^ 293TRSID-Dual-Tet-miR373 cells stably transfected with pFKis373F’Kid or with pFKis373F’Kid18 were seeded in sextuplicates per sample and allowed to adhere for 24 h, before adding doxycycline (1 μg/ml) to half of the wells for each sample. Fresh medium (containing doxycycline where necessary) was added every 3 days, taking care of collecting and reseeding back all floating cells for each sample during medium exchange. Cell were collected 6 and 9 days after addition of doxycycline, stained with Cy5-conjugated Annexin-V (Source Bioscience Autogen) following manufacturer’s instructions, and analysed by flow cytometry. Experiments were carried out in triplicates, and data plotted as the percentage of annexin-V-positive cells in each sample.

### Statistical analysis

To evaluate the statistical significance of the differences between relative values Kid/Kid18 we first applied the Shapiro-Wilt test, to determine the parametric or nonparametric distribution of our data. Once this was established, the T-Student or the Mann-Whitney test were applied to determine the statistical significance with a 90% (Figure 2E) or 95% (rest of figures) confident interval.

## Supporting information

Supplemental Methods, Table, figures and references

## Acknowledgments

We thank M. Murray and N. Coleman for sharing GCT cell lines PA-1 and 2102Ep, as well as M. Ausserlechner and R. Agami for sharing 293TRSID cells and plasmid pMSCVBlastmiRVec373, respectively. We also thank Ch. Agu and A. Arnáiz-Vivas for helpful discussions. This work was supported by funds from the Medical Research Council UK (Programme Grant MC_U105365008 to G. de la Cueva-Méndez) and from the Andalusian Ministry of Health.

## Author contributions

G.C-M. conceived the project. A.T., B.P. and G.C-M. designed the experiments. A.T., C.B-R., M.A.P, and B.P. performed the experiments and, together with M.G-B. and G.C-M, analyzed the data. A.T, B.P. and G.C-M. wrote the manuscript.

