## Supplemental Methods, Table, figures and references for "Targeted cancer cell killing by highly selective miRNA-triggered activation of a prokaryotic toxin-antitoxin system"

^3^Nanobioengineering of Smart Therapeutic and Diagnostic Systems Laboratory, Area of Oncology and Oncohematology. Instituto de Investigación Biomédica de Málaga-IBIMA.

* Co-corresponding authors, to whom correspondence should be addressed.

**Supplemental information**

Contains: i) a scheme of the steps followed to make the plasmids and cell lines described in this work, and detailed methods for their construction, as well as for cell transfection and cell cultures and statistical analysis and ii) a table describing oligonucleotides used in this work, iii) images of coomassie-stained gels identical to those used in western blots shown in Figs. 2D, 3 and 4B, iv) images of 293T-DualTet-miR373/pFKis373F’Kid cultures, before and three days after addition of doxycycline, v) histograms and box plots representing changes in net metabolic activities, and both total and dead cells numbers over time in 293T-DualTet-miR373-FKis_373_F’Kid cultures grown in the absence of doxycycline and transfected with 10 pmols of miR373, miR372, miR518f and miR302d.

**Material and Methods for Supporting information.**

**Plasmid construction.** A KpnI-hsa-miR373-BamHI DNA insert was produced by PCR with oligos K-miR373-ss and miR373-B-as, using plasmid pMSCVBlastmiRVec373 as template.^20^ The resulting DNA fragment (i.e. the endogenous stem-loop hsa-pri-miR373) was inserted between KpnI and BamHI in pTRE-Hyg (Clontech) to produce pTet-miR373. Construction of plasmids pFKis_373_F’Kid, pFKisF’Kid and their Kid18 variants was carried out in several steps. First, we used complementary DNA oligonucleotides to produce a dsDNA fragment bearing restrictions sites XmaI-AscI-NotI-SfiI-SacI-PmeI, and subcloned it between ZraI and SapI in pUC18. The resulting plasmid (pUC-Shuttle; Figure 1a in supporting information) was then used as the recipient vector for four different synthetic transcriptional units (Figure 4b in supporting information). In the first one, a human-codon-optimized *kis* gene fused to the coding sequence of a FLAG epitope was flanked by the human UbiC promoter (PrUbiC) and the SV40 early polyadenylation sequence (pA1). This transcriptional unit also included a consensus Kozak sequence immediately upstream of the start

codon of FLAG-kis, as well as single XbaI and BclI restriction sites immediately downstream of its stop codon, and flanking XmaI (5’ end) and NotI (3’ end) restriction sites, that were used to insert the cassette in the shuttle vector to produce pUC-shuttle-PrUbCFKis. To make the second and third transcriptional units, a human-codon-optimized *kid* (or its inactive variant; *kid18*) gene fused to the coding sequence of a FLAG epitope was flanked by a synthetic human promoter (PrJetwt)^1^ and the bovine growth hormone polyadenylation signal (pA2). In these transcriptional units the FLAG coding sequence was silently diverged from that fused to the *kis* transcriptional unit, to minimize the chances that deleterious homologous recombination could occur in plasmids bearing both units. FLAG-kid (and kid18) were also preceded by a consensus Kozak sequence, and the whole cassette was flanked by NotI (5’ end) and SfiI (3’ end) sites, which were used to insert it into the shuttle vector to produce pUC-shuttle-PrJetwtF’Kid. To form the last transcriptional unit, a blasticidin-resistant gene preceded by a consensus Kozak sequence was flanked by the murine phosphoglycerate kinase promoter (PrPGK) and a synthetic polyadenylation signal (pA3). The whole cassette was flanked by SfiI (5’ end) and PmeI (3’ end) sites, which were used to insert it into the shuttle vector to produce pUC-shuttle-PrPGK-BlastR. To produce miRts-bearing variants of pUC-shuttle-PrUbCFKis we annealed oligo pairs X-373ts-B-ss/B-373ts-X-as, X-372ts-B-ss/B-372ts-X-as, X-518fts-B-ss/B-518fts-X-as, X-302dts-B-ss/B-302dts-X-as, and X-artifts-B-ss/B-artifts-X-as (Figure 1 in supporting information), and subcloned them between XbaI and BclI in pUC-shuttle-PrUbC-Fkis , which produced plasmids pUC-shuttle-PrUbC-Fkis-miRts373, pUC-shuttle-PrUbC-Fkis-miRts372, pUC-shuttle-PrUbC-Fkis-miRts518f, pUC-shuttle-PrUbC-Fkis-miRts302d and pUC-shuttle-PrUbC-Fkis-miRtsartif, respectively (Figures 1C and 1D in supporting information). XmaI-NotI inserts from the plasmids above (bearing the *kis* gene) were inserted between the same sites in pUC-shuttle-PrPGK-BlastR, and the resulting plasmids were used as recipient vectors for F’kid and F’kid18 transcriptional units, which were transferred from pUC-shuttle-PrJetwtF’Kid and pUC-shuttle-PrJetwtF’Kid18 vectors using restriction with NotI and SfiI. This produced pUCFKisF’Kid (or Kid18) and all their miRTs variants (i.e. pUCFKis_373_F’Kid (or Kid18), and so on).

**Cell line construction, transfections and culture conditions.** Cells were grown at 37° C and 5% CO_2_, splitting cultures at 90% confluency. To produce 293TRSID-Tet-miR373, we stably transfected 293TRSID cells (a puromycin-resistant HEK293 derivative stably expressing Dual-Tet regulators^2,3^) with pTRE-miR373, using Effectene (Qiagen), and following manufacturer´s instructions. These cells were grown in DMEM and 10% FCS, and selection of stable transfectants was carried out in the presence of hygromycin (100 μg/ml) and the absence of doxycycline. Suitable clones were identified by comparing miR373 expression in hygromycin resistant cells, both before and after being exposed to 1μg/ml of doxycycline for 24h. The resulting 293TRSID-Dual-Tet-miR373 clones were then transfected with NotI-linearized pFKis_373_F’Kid or pFKis_373_F’Kid18, and stable transfectants were selected in DMEM, 10% FCS and 10 μg/ml blasticidin. Selection was carried out in doxycycline-free conditions to avoid inducing miR373 expression and any deleterious effect that the concomitant inhibition of FKis expression could have on cell viability, thus minimizing the chances of selecting clones expressing inactive F’Kid mutants. Suitable clones were identified by examining the expression levels of FKis and F’Kid in the absence of doxycycline, and the molecular and cellular phenotypic consequences of growing cells in the presence of 1μg/ml of doxycycline. For the analysis of the miRNA target specificity we transiently transfected 10^6^ HEK293T cells with 0.5 μg of pFKisF’Kid18, pFKis_373_F’Kid18, pFKis_372_F’Kid18, pFKis_302d_F’Kid18, pFKis_518f_F’Kid18 or pFKis_artif_F’Kid18, alone or mixed with 10 pmol of each miRNA mimic under study (Ambion), using Lipofectamine (Invitrogen) and growing cells in DMEM plus 10% FCS. PA-1 cells were grown in EMEM, plus Earles basal salt solution (ATCC) and 10% HI FBS. 2102Ep cells were grown in DMEM, 10% USA-HI FCS and glutamine. For transfection of these cells we typically used 5 μg of a 10:1 (mol/mol) mixture of plasmids pFKis_373_F’Kid, pFKis_373_F’Kid18, pFKisF’Kid, or pFKisF’Kid18 and an EGFP-reporter plasmid (pMaxEGFP; Lonza). This mixture was used to transfect 2x10^6^ cells, using Buffer V and nucleofection programs X-01 (PA-1 cells) or A-20 (2102Ep cells) in an Amaxa Nucleofector device, and following instructions supplied by the manufacturer (Lonza).

For the experiment in supplementary Fig. 4, 25x10^4^ 293TRSID-Dual-Tet-miR373 cells stably transfected with pFKis373F’Kid were nucleofected with 10pmol of the indicated miRNA and 0.1 pmol of a reporter pmaxGFP plasmid using SF Amaxa Nucleofection solution and CM-130 program in a LONZA Nucleofector 4D equipment. Conditions were optimized following manufacturer’s instructions to obtain 80% nucleofection efficiency (monitored by GFP expression at 32 hrs post nucleofection). Nucleofected cells were seeded in triplicate for each sample and time point to be analyzed (16, 32, 64, and 82 h), using 96-well plates and cultured in the absence of doxycycline. Cells were incubated with 2.5 μg/ml Hoechst33342 nuclei stain and 0.5 μg/ml of Propidium iodide for 30 minutes and then analyzed in a Perkin Elmer Operetta High Content Microscope at the indicated timepoints. The net metabolic activity in our samples was determined at the indicated time points using the Cell Counting Kit 8 (CCK-8; Dojinjo), a colorimetric assay that quantifies the amount of NADH and NADPH (a surrogate for metabolic activity, and therefore for cell viability and cell numbers) produced by cultured cells, following manufacturer´s instructions. Experiments were carried out in triplicates, and data plotted as fold change in metabolic activity (Fig. 4a), total cell numbers (Fig. 4c), or dead cell numbers (Fig. 4c) with respect to those observed in control cells transfected with the reporter plasmid alone. Box plots show all measurements per experiment without normalization against control cells.

**Statistical analysis**. For the evaluation of statistical significance of the differences between relative or absolute values of two experimental conditions we first applied the Shapiro-Wilt test, to determine the parametric or nonparametric distribution of our data. Once this was established, the T-Student or the Mann-Whitney test were applied to determine the statistical significance with a 90% or 95% confident interval

**Supplemental information References**

1. Tornøe, J., Kusk, P., Johansen, T.E., Jensen, P.R. (2002). Generation of a synthetic mammalian promoter library by modification of sequences spacing transcription factor binding sites. *Gene 297*, 21–32.

2. Agu, C.A., Klein, R., Schwab, S., König-Schuster, M., Kodajova, P., Ausserlechner, M., Binishofer, B., Bläsi, U., Salmons, B., Günzburg, W.H., Hohenadl, C. (2006) The cytotoxic activity of the bacteriophage lambda-holin protein reduces tumour growth rates in mammary cancer cell xenograft models. *J. Gene. Med. 8*, 229-241.

3. Ausserlechner, M.J., Obexer, P., Deutschmann, A., Geiger, K., Kofler, R. (2006) A retroviral expression system based on tetracycline-regulated tricistronic transactivator/repressor vectors for functional analyses of antiproliferative and toxic genes. Mol Cancer Ther. 5, 1927-1934.

**Supplemental Figures**

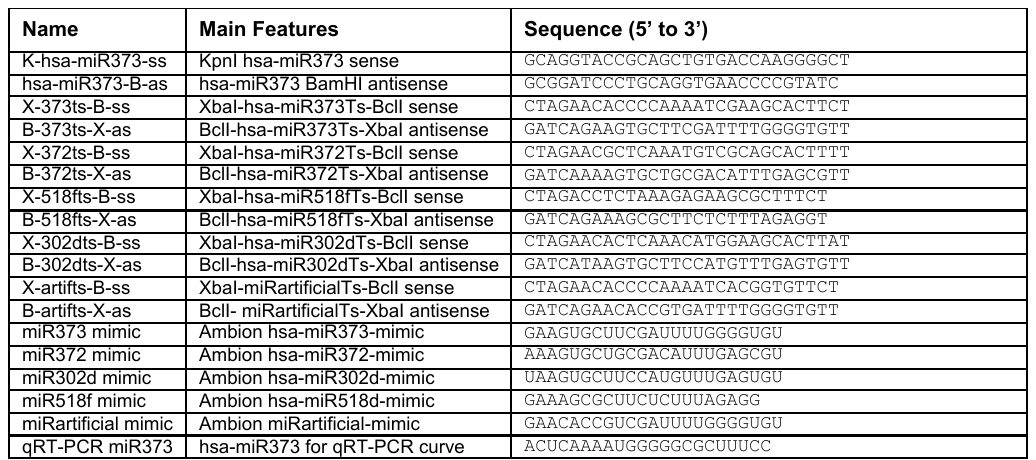

**Supporting Information Table 1.** Name, sequence and brief description of DNA and RNA oligonucleotides used in this work

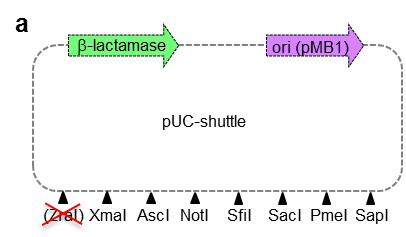

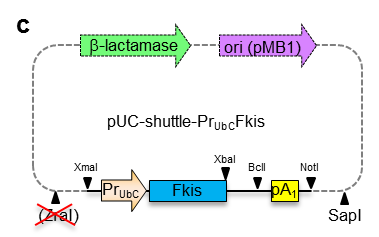

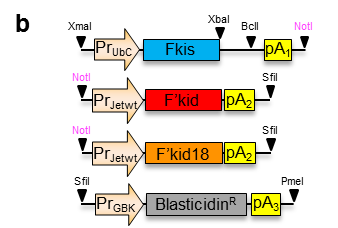

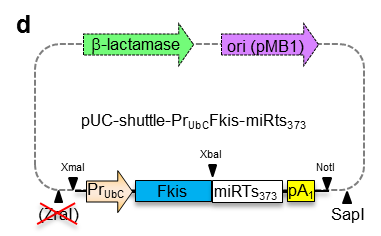

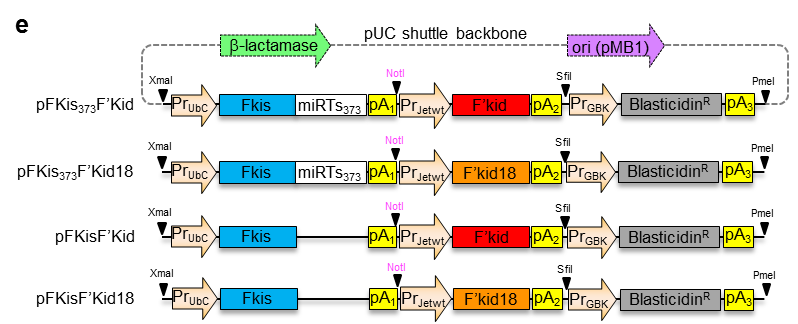

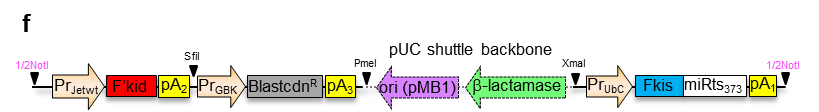

**Supporting Information Figure 1.** (a) pUC-shuttle vector used in the construction of the *kis-* and *kid-*expressing plasmids used in this work. (b) Scheme of Fkis-, F’kid-, F’kid18- and blasticidin resistance-transcriptional cassettes used in this work, showing key restriction sites used in cloning steps and in genomic integration of the final plasmids. (c) Scheme of the pUC-shuttle-Pr_Ubc_-Fkis and of a derivative plasmid bearing a target site for hsa-miR373 immediately downstream of the stop codon of *kis* (d), to illustrate how miRNA target sites are inserted in the plasmid above. (e) Scheme of all vectors produced in this work by sequential cloning of the transcriptional units shown in (b) in the plasmids shown in (c) and (d). (f) Disposition of Fkis-, F’kid-, F’kid18- and blasticidin resistance-transcriptional cassettes from plasmids shown in € once they are integrated in the genome of human cells after being linearized with NotI. Crossed red lines denote restriction sites that were inactivated during plasmid construction.

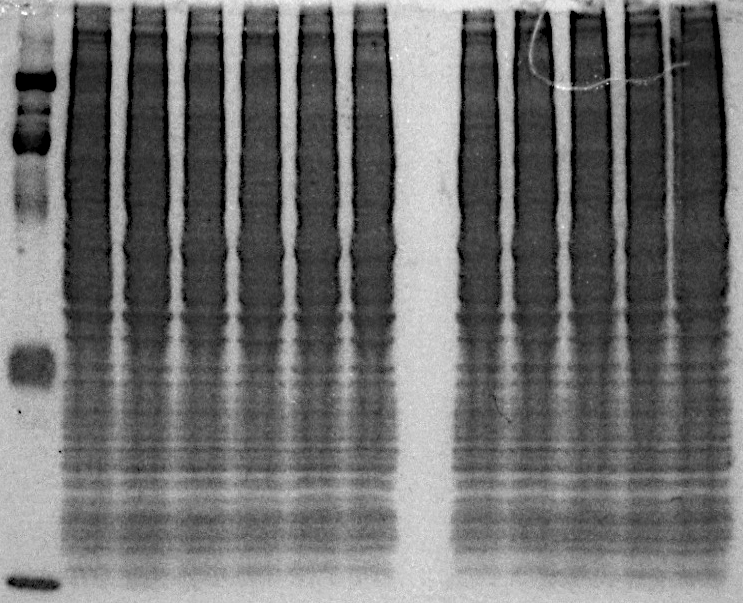

293T-DualTet-miR373/pFKis_373_F’Kid18

Ctrl

0

24

48

72

96

0

24

48

72

96

M

(hours induction miR373 expression)

Coomassie stain of gel in Fig. 2d

**a**

Coomassie stain of gel in Fig. 3a

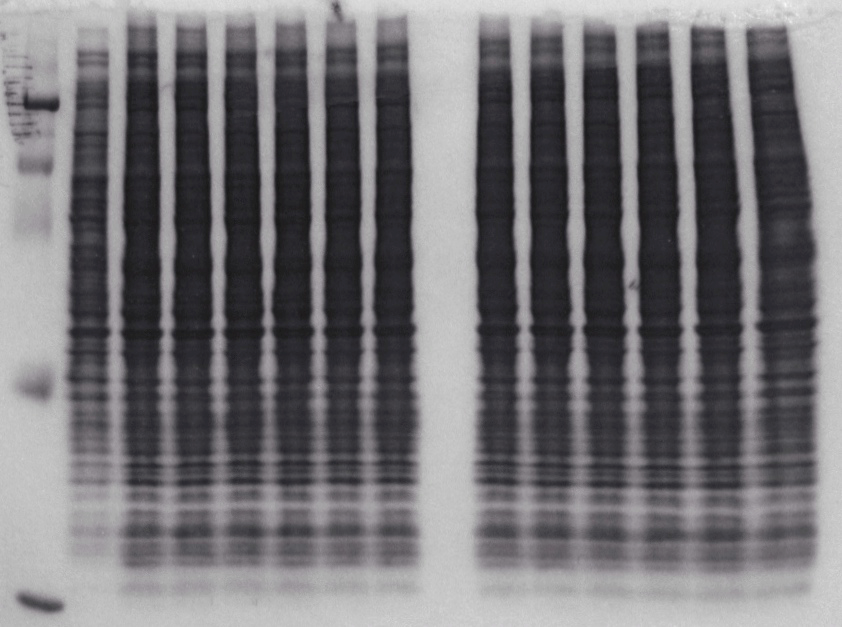

Ctrl

No miRNA

miR373

miR372

miR302d

miR518f

miRartif

No miRNA

miR373

miR372

miR302d

miR518f

miRartif

MWM

**b**

No miRNA

No miRNA

No miRNA

No miRNA

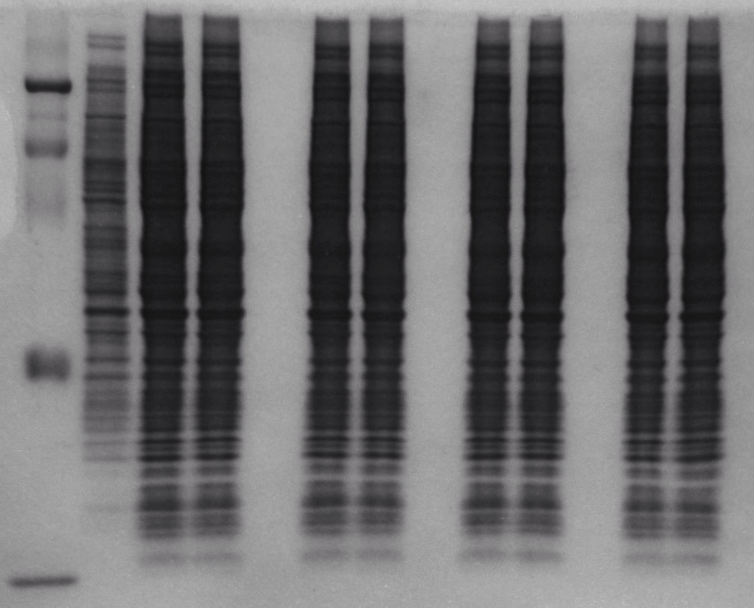

Ctrl

miR372

miR302d

miR518f

miRartif

FKis_miRTs_F’Kid18

miRTs372

miRTs302d

miRTs518f

miRTsartif

MWM

**c**

Coomassie stain of gel in Fig. 3b

**Supporting Information Figure 2**. Coomassie stain gels identical to those use for western blots in a) Fig 2d, b) Fig 3a and c) Fig 3b

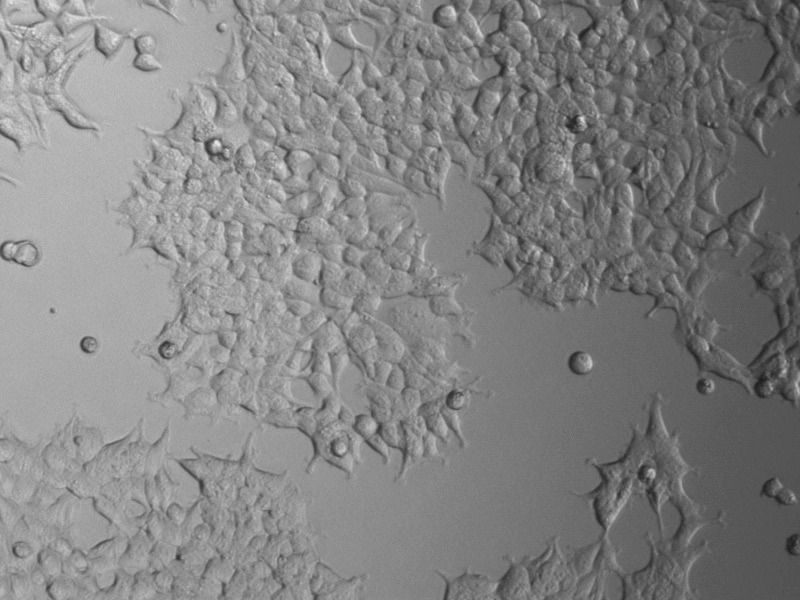

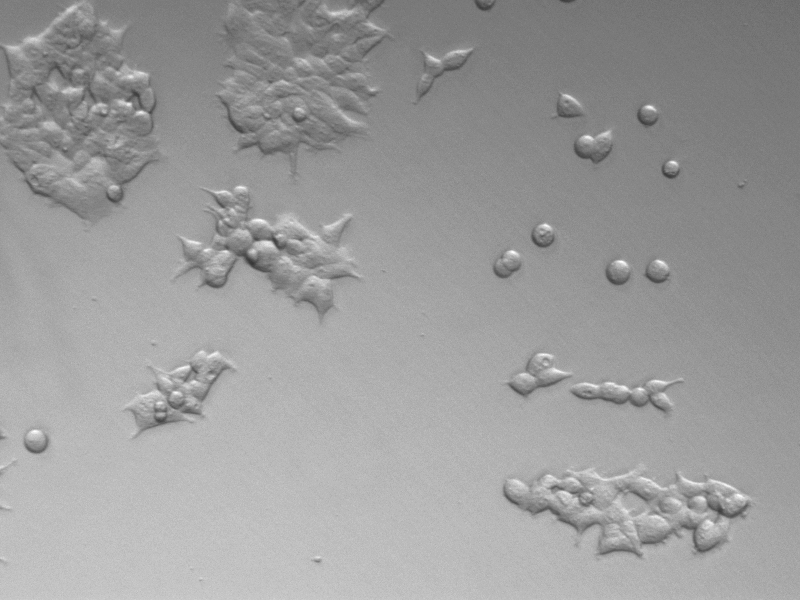

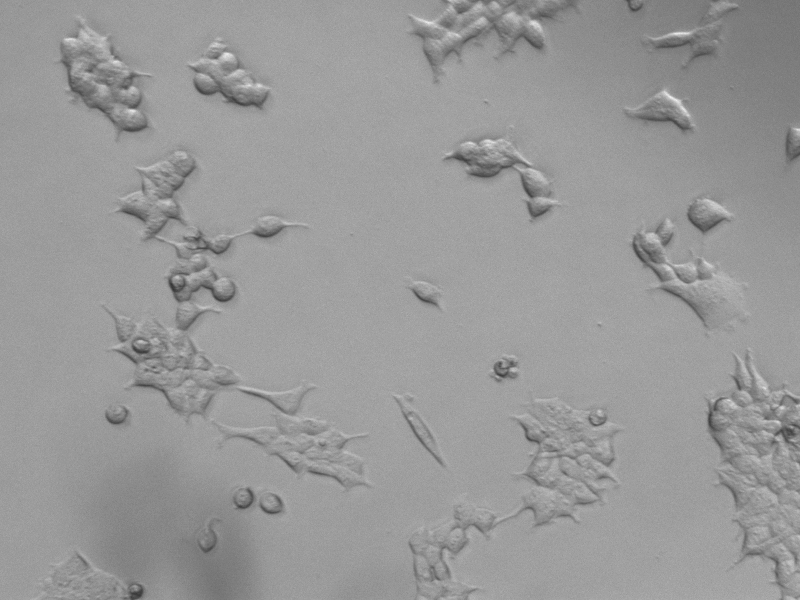

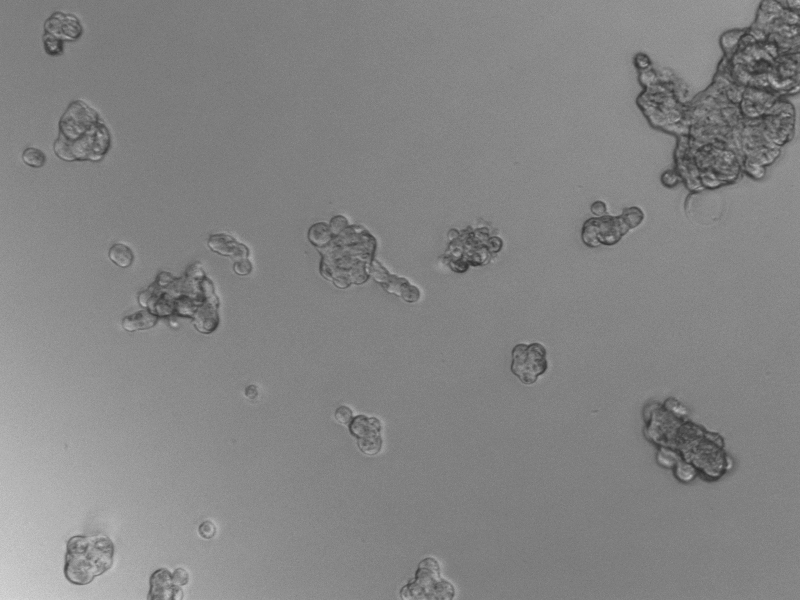

293T-DualTet-miR373/pFKis_373_F’Kid

Day 0 / no Dox

Day 3 / no Dox

Day 0/ Dox

Day 3/ Dox

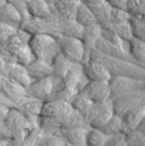

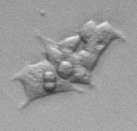

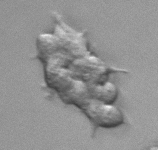

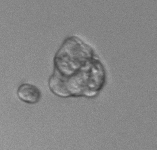

Day 0 / no Dox

Day 3 / no Dox

Day 0/ Dox

Day 3/ Dox

**Supporting Information Figure 3**. Microscopy image of the 293TRSID-Dual-Tet-miR373-pFKis373F’Kid cell line cultured in the absence (no Dox) and in the presence of 1ug/ml of Doxycycline for 3 days (Dox). Scale bar of the top image is 12 um. Bottom image show amplify regions displayed in top images

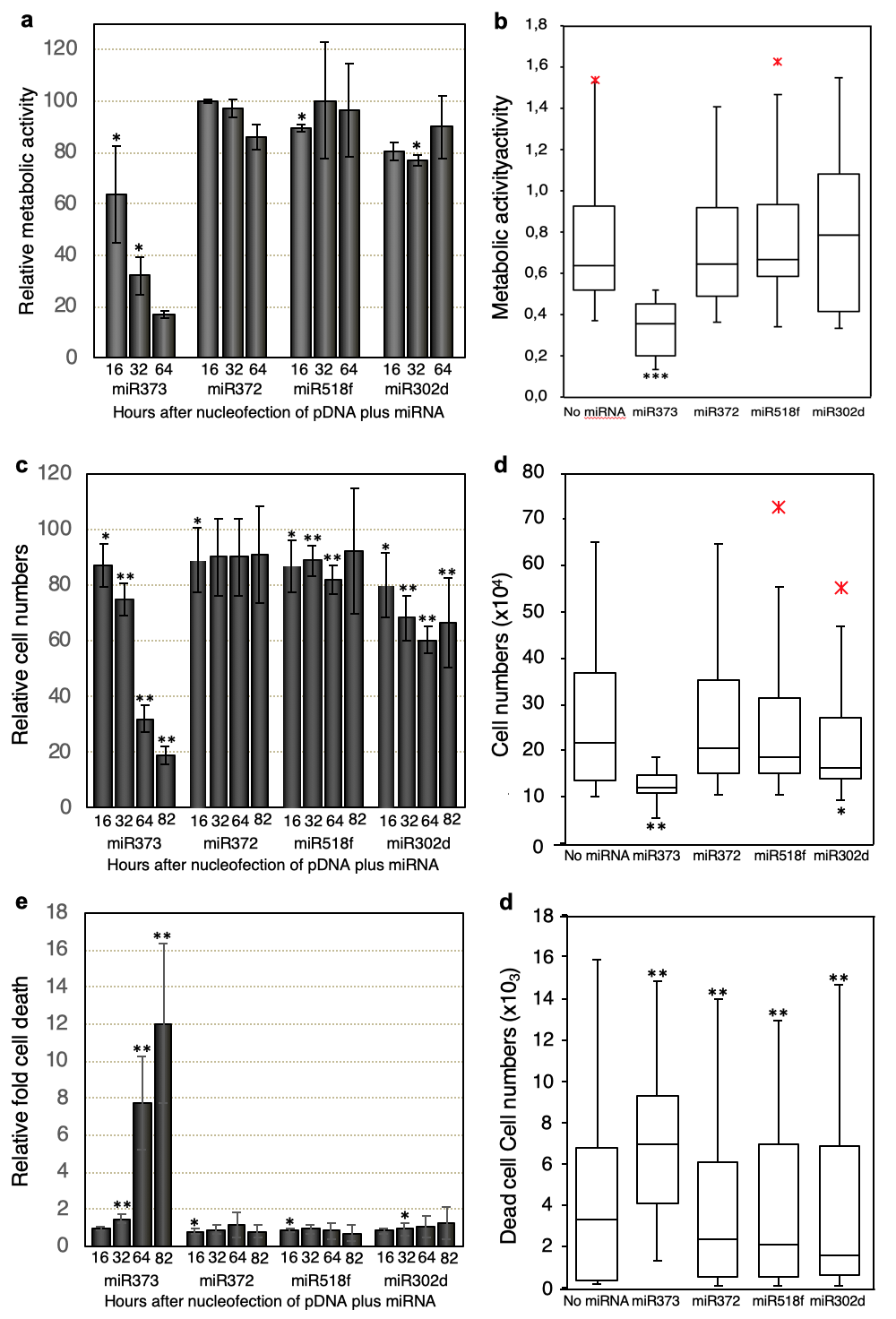

**f**

**Supporting Information Figure 4**. Fold change of the a) average net metabolic activity, c) total number of cells, and e) total number of dead cells in 293TRSID-Dual-Tet-miR373-pFKis373F’Kid cultures measured at the indicated time points after co-transfection with 10pmol of the indicated miRNA and 1 pmol of a pmaxGFP reporter plasmid. Fold change is relative to same cells transfected with the reporter plasmid alone. Panels b), d), and f) are box plots of all measurements, not releative to control cells, which are plotted independently, for all experiments (n=3) shown in panels a), c) and e). * and ** denote a statistically significant difference at 90% and 95% confidence interval, respectively, between the control reporter plasmid alone and those samples transfected with both the reporter plasmid and the indicated miRNAs. Experiments were repeated three times, and all measurements were carried out in triplicates.
